# BCL7 proteins, metazoan-specific subunits of the mammalian SWI/SNF complex, bind the nucleosome core particle

**DOI:** 10.1101/2023.03.17.532992

**Authors:** Dana Diaz, Asgar Abbas Kazrani, Franck Martin, Julie Lafouge, Stéphanie Siebert, Catherine Birck, Alexandre Blais, Elisa Bergamin

## Abstract

BCL7 proteins are among the most recently identified subunits of the mammalian SWI/SNF (mSWI/SNF) chromatin remodeling complex and are absent from the unicellular version of the complex. Mutations in BCL7 proteins are associated with different kind of cancers including blood malignancies. The information on the molecular function and on the structure of BCL7 proteins is to date very limited. Here we report that BCL7 proteins directly bind the nucleosome core particle (NCP) and free DNA with high affinity. We demonstrate that BCL7 proteins form defined complexes with the NCP and we identify the conserved N-terminal part of BCL7 proteins as sufficient to nucleosome binding. We further characterize the impact of BCL7 protein mutations reported in cancer patients on NCP binding and show that the R11S driver mutation reduces the affinity for the nucleosome. Our findings clarify the molecular function of BCL7 proteins and help rationalize the impact of cancer-associated mutations.

## INTRODUCTION

At the most fundamental level, eukaryotic DNA is packaged into chromatin by making nearly two turns around an octamer of histone proteins, forming nucleosomes. Several multi-protein, chromatin-modifying complexes contain at their core an ATPase subunit that remodels chromatin: replacement of core histones with histone variants that influence DNA accessibility; sliding, addition or eviction of nucleosomes from the chromatin fibre to modulate the compaction of DNA or expose certain regions^1,2^. Deregulation of this process can severely impact gene expression patterns and genome integrity. Their importance is such that chromatin remodelling protein mutations are strongly associated to several diseases, including cancer. Unfortunately, we currently do not understand well enough the molecular details of their mode of action to be able to translate this into improved cancer treatment.

All eukaryotes have multiple ATP-dependent chromatin remodelling complexes, which tend to be highly conserved, from simple yeasts to metazoans, testifying to their importance. Their ATPases all have homology to the *S. cerevisiae* enzyme SNF2 and are classified into sub-families based on the amino acid sequence of the ATPase: SNF2, ISWI, CHD, and INO80. The complexes accomplish their remodelling function by antagonizing the strong interaction between the histone octamer and DNA, to alter chromatin structure. In addition to their ATPase subunit, these complexes contain more associated proteins that contribute to their function. The mammalian switch/sucrose non-fermentable (mSWI/SNF) complex appears as a ∼ 2MDa complex and is composed of at least 11 subunits, coming from the product of 29 genes and multiple paralogs generating wide diversity in its composition. mSWI/SNF complexes exist in three forms: canonical BAF (BAF or cBAF) complexes, polybromo-associated BAF (PBAF) complexes, and non-canonical BAF (ncBAF) complexes, each distinguished by the incorporation of distinct accessory subunits. Each complex comprises a set of evolutionary conserved ‘core and enzymatic’ subunits, but also ‘auxiliary’ subunits present only in animals thought to reflect increasing biological complexity. All complexes contain a conserved ATPase subunit, either SMARCA4 (BRG1) or SMARCA2 (BRM), that catalyses the hydrolysis of ATP. The advent of sensitive and large-scale proteomics methods has shed new light on the composition of the human SWI/SNF. Studies focused on identifying mSWI/SNF components^3,4^, as well as others determining the composition of most nuclear multiprotein complexes^5,6,7^ have revealed the identity of previously unsuspected mSWI/SNF subunits such as BCL7A, BCL7B, BCL7C, BRD7, BRD9, SS18, SS18L1, GLTSCR1. Without exception, these proteins do not exist in yeast, and only participate in the metazoan complexes. The molecular function of most accessory subunits and their role within mSWI/SNF remain incompletely defined ^8,9,10,11,12,13,14^. It has been reported that BCL7 proteins accumulate at sites of DNA damage, but the role they may play in DNA damage repair (DDR) is unknown ^15^.

BCL7A is labelled as “Cancer Genes” by the COSMIC database as it is often mutated in a variety of cancers and established tumor suppressors^16,17^. Despite this, the structure, the molecular details of interaction and the function of BCL7 subunits within mSWI/SNF are poorly defined at best. The topological information currently available is limited to the observation that BCL7 proteins make contacts with BRG1, SMARCB1, DPF2, actin and histone H2B^18,19^. Recent studies have reported the cryo-EM structure of the SWI/SNF complex. Unfortunately, they do not inform us on BCL7 proteins: two studies concerned the yeast SWI/SNF complex and BCL7 are absent in this species^20,21^; in studies of reconstituted mSWI/SNF, BCL7 proteins were either omitted^22,23^ or not detectable^19^; and studies with the endogenous human complex had overall poor resolution and does not provide molecular details on BCL7 proteins^24^.

Interestingly, although mSWI/SNF is an NCP-displacing multi protein complex, until recently only SMARCB1 and the ATPase BRG1^25,26^, had been reported as proteins carrying clearly mapped NCP-binding domains. The interaction of PBAF with the nucleosome has been further clarified with the identification of more subunits coming in contact with the nucleosome^19^, but much remains to be discovered about the interactions between mSWI/SNF and its substrate.

We have undertaken work to address this knowledge gap by investigating the role of the BCL7 proteins, since understanding how BCL7 proteins interact with the nucleosome is essential to understand the function of these enigmatic proteins in health and disease. We used biochemical and biophysical approaches and obtained evidence that BCL7 proteins directly bind DNA and the NCP and can form a stable complex with it. Truncation experiments show that the N-terminal part of the protein is sufficient for NCP binding. We find that single amino acids substitutions within the conserved N-terminal part of BCL7A, which are frequent in cancer patients, impair NCP binding.

## EXPERIMENTAL PROCEDURES

### Protein expression and purification

The cDNA encoding full length human BCL7A (isoform Q4VC05-2, 231 amino acids) and BCL7A (1-100), full length BCL7B and BCL7C, were subcloned into the parallel vector pGST2 that includes a TEV-cleavable, N-terminal glutathione sulfotransferase (GST) tag. Each vector encoding BCL7 proteins was transformed into *E. coli* strain Rosetta (DE3), and cultures were grown in Terrific Broth media at 37° C to an OD_600_ of 0.6. Protein expression was induced by the addition of IPTG (0.2 mM) for 16 hours at 19° C. Cells were harvested, resuspended in lysis buffer (50 mM Tris [pH 8.0], 300 mM NaCl, 0.1% [v/v] Triton X-100, 5 mM 2-mercaptoethanol), lysed by sonication and clarified by centrifugation. The supernatant was collected and applied onto glutathione Sepharose beads for 2 hours at 4°C and washed extensively with lysis buffer. GST-BCL7 was TEV cleaved on beads for 16 hours at 4°C. The untagged protein was further purified by heparin affinity chromatography followed by size exclusion chromatography (Superdex 200) preequilibrated in 20mM Tris [pH 8.0], 150mM NaCl and 1 mM DTT. BRG1 was overexpressed in Sf9 cells and purified as previously reported ^27^.

### Nucleosomal DNA and large-scale nucleosome reconstitution

147-bp palindromic DNA fragments derived from the strong positioning 601 DNA sequence^28^ was used. To produce sufficient quantity of nucleosomes for biochemical studies, a large-scale protocol was employed. The DNA was prepared in milligram quantities using a plasmid with multiple concatenated copies of the 601 sequence, followed by restriction digestion of the individual units and size selection, as reported previously^29^. Core histones were produced by co-expression in *E. coli* and were purified in native form following an established protocol^30^. The two components were assembled into NCPs by over-night dialysis to reduce NaCl from 2M to 0.25M. NCPs were heat-shifted at 37 ° C for 120 minutes. Proper assembly was assessed on non-denaturing polyacrylamide gels (5% acrylamide 37.5:1 acrylamide:bis-acrylamide ratio, 0.25X TGE, 0.03% v/v NP-40) pre-run for 1 hour at 4 ° C and run for 1 hour at 120 volts on Mini-Protean 3 gels (Bio-Rad).

### Electrophoretic mobility shift assays (EMSA)

Fluorescent nucleosomes were assembled using the salt dilution method with recombinant histones and Cy5-labelled 601 DNA produced by PCR. The labelled probes were incubated at 10 nM in 30 μl binding reactions containing 50 mM Tris-HCl pH 8.5, 20 mM KCl, 10 mM β-mercaptoethanol, 10% (v/v) glycerol. Purified proteins were omitted or added to the binding reactions at six varying concentrations ranging from 75 to 1800 nM. The concentrations were chosen after preliminary tests with a broader concentration range, and selected so that in general there was a large molar excess of protein over ligand (total protein is a reasonable approximation of free protein and binding reactions occurred in the “binding regime”)^31^. Reactions were incubated on ice for 30 minutes and then loaded on non-denaturing 5% acrylamide gels (60:1 ratio of acrylamide to bis-acrylamide) containing 10% glycerol, using 0.5X TGE (40 mM Tris-HCl, 45 mM boric acid and 1 mM EDTA) as running buffer.

For EMSA with short oligonucleotide probes, DNA oligos were chemically synthesized with a Cy5 dye molecule attached to the 5’ end of the top strand. Oligos were annealed to make double-stranded probes by heating and slowly cooling. Probes were used in EMSA as indicated above. Oligonucleotide sequences are provided in Table S1.

For EMSA with plasmid DNA, the reactions were assembled the same way as described above but using a broader range of BCL7A concentrations. Reactions were run on acrylamide gels and stained with ethidium bromide before imaging.

### Calculation of apparent dissociation constant (Kd)

After electrophoresis, gels were immediately scanned using the Typhoon Phosphorimager, at 10 micrometer resolution, adjusting the PMT value in order to obtain the strongest signal without saturating any pixel. Image files were saved in 16-bit.gel format and imported into the ImageQuant TL 10.1 software (GE Healthcare). Free NCP signal was quantified in boxes of equal area as relative intensities, and background was subtracted using the “rubber band” method. The free NCP values in each lane were used, expressed as fraction of free NCP in the absence of protein, and fitted to a quadratic binding equation^31^, fitting the fraction of free probe (labelled NCP or DNA) as a function of the micromolar concentration of protein, and total ligand concentration in the assay (0.01 µM). Fitting was performed using the nlsLM function from the ‘minpack.lm’ package in R. The initial value for Kd was set to 0.2. Using an initial value 10 times higher or 90% lower did not affect the results. Multiple replicates of each experiment were performed. For each replicate, the residuals sum of squares from measured *versus* fitted values was used to perform a goodness-of-fit analysis; this returned Chi-squared p values lower than 1E-4 in every case. The final reported values are the arithmetic mean ± standard error of replicates. Student’s t tests (unpaired, two-tailed, equal variance) were used to determine statistical significance.

### Microscale Thermophoresis (MST) experiments

MST experiments were performed on a Monolith NT.115 (NanoTemper, LLC, Munich, Germany) using premium coated capillaries. All data sets were collected at 25° C. Reactions were performed in buffer consisting of 20 mM HEPES pH 8.0, 150 mM NaCl, 0.05% Tween-20. BCL7A was prepared using 1:1 serial dilution of the protein into the buffer with the highest concentration of 30 µM. Sixteen 10-μL samples were thus prepared ranging in concentration from 30 μM to ∼1 nM. The association was initiated by the addition of an equal volume of 40 nM Cy5 labelled NCP. The mixtures were incubated in the dark at room temperature for 10 minutes before being loaded into capillary tubes and inserted into the apparatus for data acquisition. The LED power was 80% and the MST power was 20%. Data analysis was performed in the Nano Temper Analysis software using a *Kd* fitting analysis on temperature jump data. Triplicates were used for data fitting. Data points at the extremes of the range were excluded from the analysis. Straight lines were fitted to the unsaturated and saturated portions of the data in triplicate using GraphPad Prism.

### Analytical ultracentrifugation

Sedimentation velocity (SV) experiments were performed at 20°C and 42,000 RPM in a Beckman Coulter ProteomeLab XL-I ultracentrifuge with an AnTi-50 eight-hole rotor. NCP was further purified on gel filtration column before SV experiment. Samples of 420 µl or 110 µl were loaded into analytical cell assemblies with 12- or 3-mm charcoal-filled Epon double-sector centerpieces and sapphire windows. The reference buffer was 20 mM Tris pH 8, 100 mM NaCl, 1 mM DTT. After temperature equilibration at 20°C for 3h, the rotor was accelerated to 42000 rpm and 300 scans were collected with absorbance optics at 280 nm and interference optics. Sedimentation data were analyzed with SEDFIT software (version 16.1c) using the continuous sedimentation coefficient distribution model c(s)^32^. Buffer density, buffer viscosity and protein partial specific volumes were calculated using SEDNTERP software. A partial specific volume of 0.65 mL/g was used for NCP or NCP+ BCL7A, and 0.7107 mL/g for BCL7A. A solvent density of 1.03 g/mL and a solvent viscosity of 1.017 cP were used for data analysis. GUSSI was used to integrate the sedimentation peaks and to produce the graphs ^33^.

### Size exclusion chromatography / multi-angle light scattering

Size exclusion chromatography combined with multi-angle light scattering (SEC-MALS) was performed with a Superose 6 (10/300 GL) column at 0.5 ml/min, 25ºC, pre-equilibrated in 20 mM Tris pH 8.0, 150 mM NaCl, 1 mM DTT, to separate the sample before performing the MALS measurements. BCL7A wild type, nucleosome alone or in presence of a 1:50 molar excess of BCL7A were individually injected in a volume of 50 µl of sample at 5 mg/ml. The molar mass for each molecule was determined with the ASTRA software (Wyatt Technologies).

### Cancer mutation analysis

The COSMIC database was queried on 2023-01-10 for samples with mutations to the BCL7A gene and the results were retrieved for plotting. The number of samples with mutation at each amino acid position, regardless of the change (missense, nonsense, insertion or deletion) are reported. For the DLBCL mutation hotspot analysis, the data from the FishHook analysis by Mlynarczyk et al. were downloaded^34^. This analysis was performed with a cohort of 101 DLBCL cases from the ICGC/TCGA Pan-Cancer Analysis of Whole Genomes Consortium^35^. As reported by the authors, a Q-Q plot of the log-transformed observed P values versus the log-transformed quantiles from the uniform distribution was prepared.

## RESULTS

### Sequence comparison and structural predictions of BCL7 proteins

The BCL7 family is composed of three proteins encoded by three different genes: BCL7A, B and C. They lack any domain of known molecular function or shared with other proteins. The longest mRNA isoform of human BCL7A comprises six exons and codes for a protein of 231 amino acids. A second mRNA isoform exists that shortens the end of exon 5 by 21 codons. According to multiple sequence alignment results of the three human BCL7 family members, the first 51 amino acids of each protein are highly homologous, while homology is more limited beyond this position (Fig. 1A and B). The sequence of each family member is conserved throughout evolution across the entire length of the proteins (Fig. S1). Sequence-based analyses using DISOPRED and AlphaFold ^36,37^ predict that the majority of the protein’s length is disordered (Fig. 1C and D).

**Figure 1.**
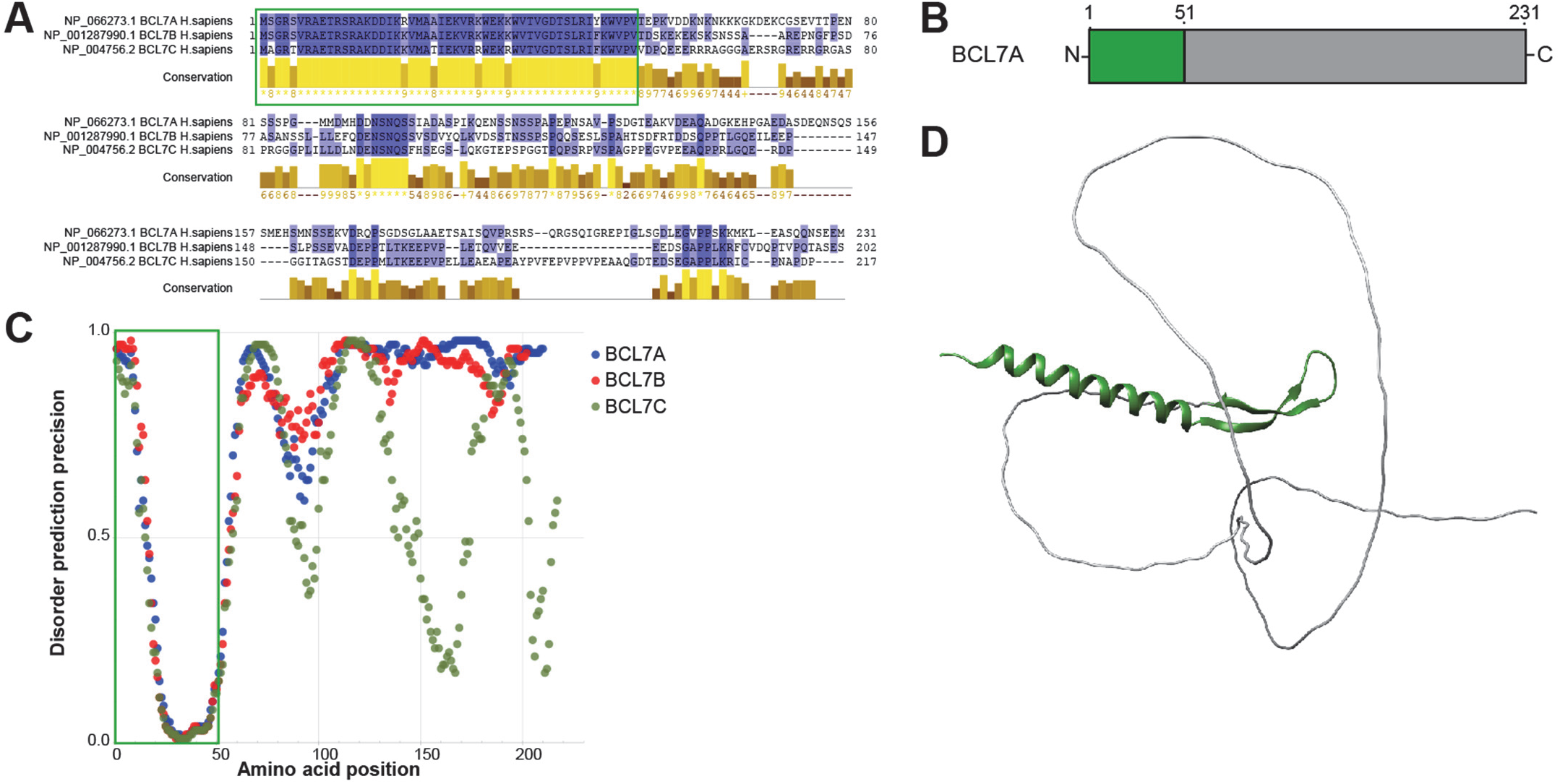
Sequence and disorder of BCL7 proteins. **A)** Multiple sequence alignment of the three human BCL7 proteins performed using Muscle in JalView. The first 51 amino acids, highly homologous between the three proteins, are highlighted in a green box. They correspond to the BCL_N domain reported in the Pfam database of protein domains (PF04714). **B)** Domain architecture of BCL7A. **C)** Disorder prediction for the three human BCL7 proteins, obtained using DISOPRED3. The region corresponding to the first 51 amino acids is shown in a green box. **D)** AlphaFold prediction of the structure of BCL7A. The N-terminal homologous region in the green box of panel A is represented in green.

### BCL7 proteins form complexes with nucleosomes and free DNA

In initial recombinant protein production experiments, it was noticed that the three BCL7 proteins tend to co-purify from *E. coli* lysates along with significant amounts of bacterial host DNA (data not shown), which suggested a possible direct DNA-binding function. Electrophoretic mobility shift assays (EMSA) confirmed this intuition. EMSA performed with recombinant human BCL7A and a variety of labelled DNA probes revealed a protein concentration-dependent gradual decrease in probe mobility indicating stable complex formation. The probes were 12 to 16 base pairs in length and varied in types of DNA ends (blunt, 5’- or 3’-protruding, Fig. 2A and 2D), base composition (only A-T or G-C base pairs, Fig. 2B), and cytosine methylation (with or without methyl-CpG, Fig. 2C), but no obvious difference in probe binding activity was observed. The ability of BCL7A to bind DNA without ends (i.e. circular plasmid) was also observed (Fig. 2D).

**Figure 2.**
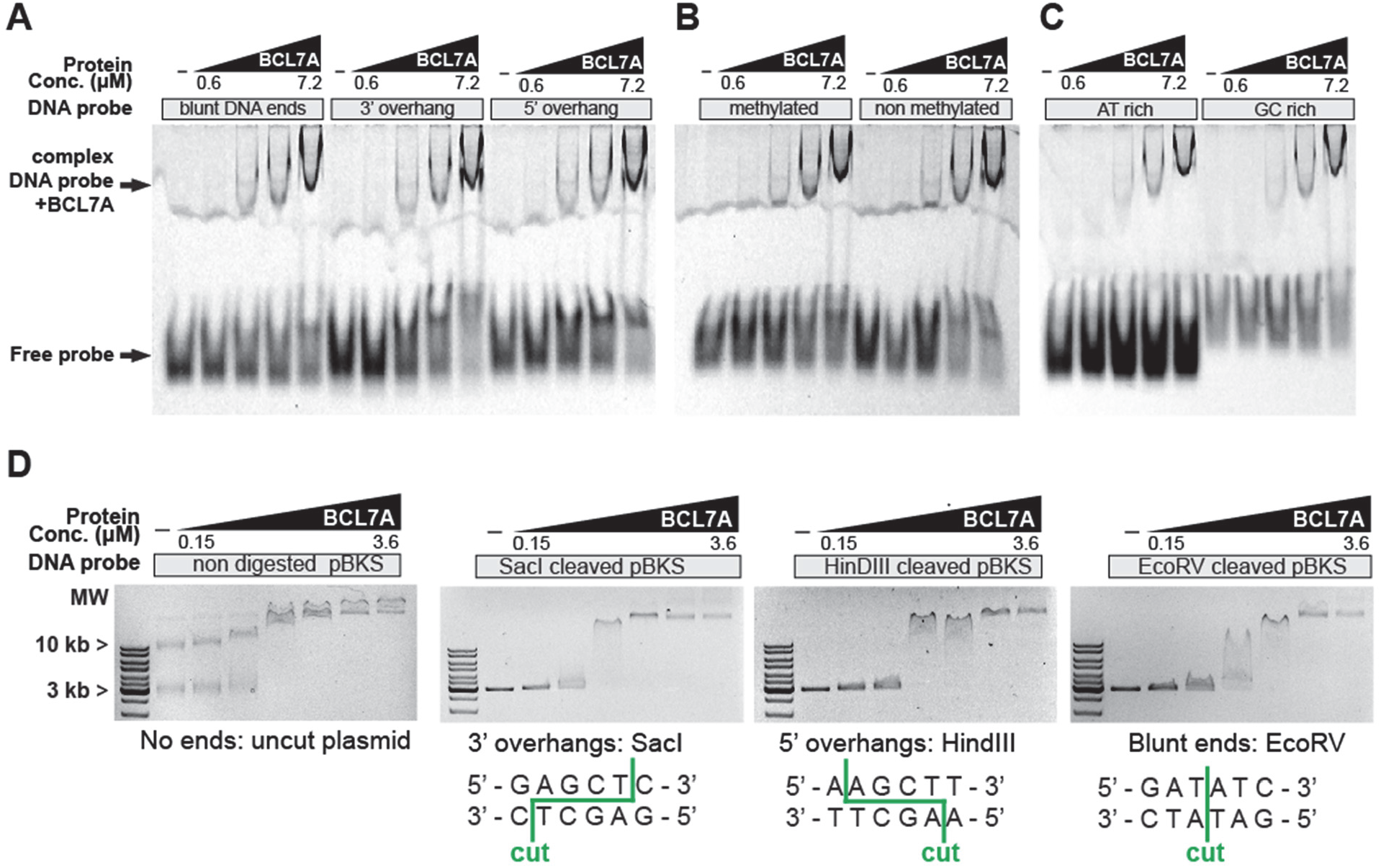
BCL7A binding preference over different DNA probes. EMSA experiments were performed with increasing amounts of BCL7A WT and a variety of DNA probes to test binding preference over different DNA ends (A), DNA methylation (B) and DNA base composition (C). BCL7A does not show any preferential binding specificity. D) On plasmid probes, BCL7A does not show any preference over the presence or absence of DNA ends, or on the nature of DNA ends. The 2.9 kb pBS-KS plasmid was used. The large molecular weight DNA species seen in the non-digested plasmid are likely contaminating *E. coli* genomic DNA, which also is shifted by the added BCL7A protein.

Because mSWI/SNF can act on nucleosomes, EMSA experiments were performed to examine binding of BCL7A to this type of probe. Nucleosomes were assembled using recombinant *X. laevis* histone octamers and fluorescently labelled 601 positioning DNA sequence ^28^. EMSA with free 601 DNA or nucleosomal probes revealed that BCL7A can bind both types of substrates (Fig. 3A). Curve fitting to explain probe electrophoretic retardation indicated that the binding affinity of BCL7A on the NCP probe has an apparent dissociation constant (Kd) of 218 nM (Fig. 3B), compared to a Kd of 237 nM on the naked DNA probe of identical sequence (Fig. 3C); the difference was not significant by Student’s t-test, indicating that the affinity of BCL7A to naked or NCP probes is similar.

**Figure 3.**
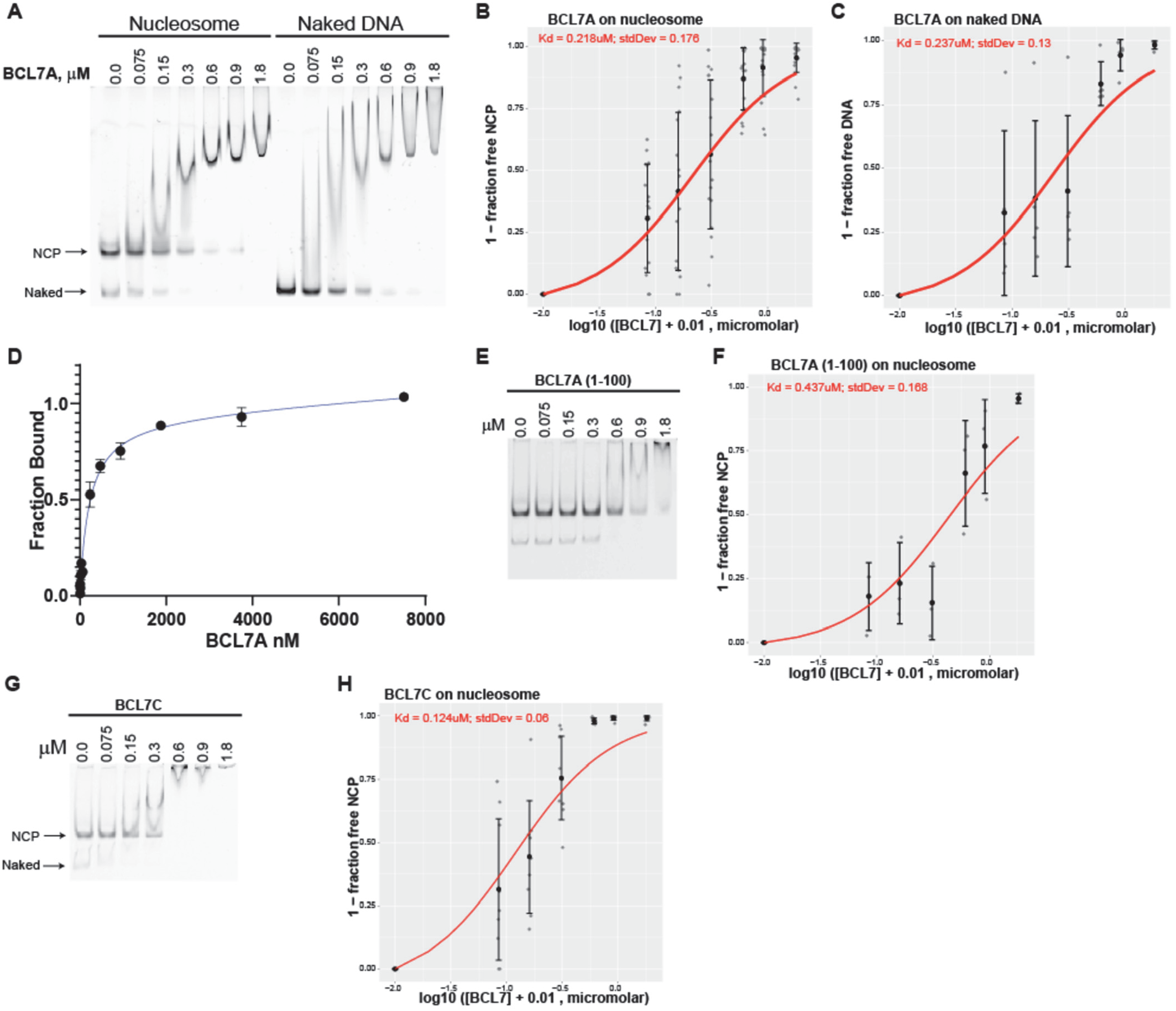
Binding of BCL7 proteins to nucleosomal DNA. **A)** EMSA experiments where constant amounts of BCL7A were incubated with 601 DNA assembled into nucleosomal core particles or left as naked DNA. **B)** Quantification of binding data for BCL7A on NCP probes. Each replicate experimental point is shown by one grey dot. The mean and standard error are shown by black brackets. The Hill equation curve fit (using the mean Kd and Hill coefficient from all replicates) is shown in red. **C)** Kd calculation for BCL7A binding to naked DNA. **D)** MST experiment using labelled NCP in constant concentration and increasing concentrations of unlabelled BCL7A protein. The points represent the average and standard errors from two independent replicates. **E and F)** Binding of BCL7A truncation mutant 1-100 to the NCP. **G and H)** Binding of BCL7C to the NCP.

To confirm the EMSA results showing high affinity binding of BCL7A to nucleosomes, multiscale thermophoresis (MST) was employed. MST experiments were performed with Cy5-labelled NCP and increasing amounts of full length BCL7A. Using this method, a binding affinity of ∼230 nM was obtained, which is close to the value of 218 nM calculated from the EMSA data (Fig.3D).

To identify which part of the BCL7A protein might be implicated in NCP binding, the EMSA experiments were repeated with the BCL7A (1-100) truncation mutant, where most of the predicted disordered C-terminal region is removed. It was found that the first 100 amino acids of BCL7A are sufficient for nucleosome binding, though the affinity is somewhat lower (Kd of 437 nM); the difference with wild-type was not statistically significant (p > 0.05 by Student’s t-test, Fig. 3E and F).

The ability of BCL7C to bind nucleosomes was also examined by EMSA, which yielded a Kd of 124 nM (Fig. 3G and H), indicating that the two proteins possess the same ability to bind to the NCP with high affinity.

### Impaired NCP binding by BCL7A protein carrying cancer-derived mutations

The three BCL7 genes have been reported to be mutated in cancer cases ^14^. Mutations of BCL7A are especially frequent, notably in haematological malignancies such as diffuse large B-cell lymphoma (DLBCL)^11,31,17^. The mutations tend to accumulate at the N-terminal region (Fig. S2A) and represent cancer driver mutations (Fig. S2B). We tested the ability of two BCL7A mutants to bind the nucleosome: R11S and P78S. It was observed that at arginine 11, mutation to serine significantly reduces the affinity of BCL7A for the nucleosome (p < 0.05, Fig. 4A). Similarly, mutation of proline 78 to a serine residue significantly decreased the affinity for the nucleosome (p < 0.05, Fig. 4B). These results suggest that the N-terminal region of BCL7A is indeed important for nucleosome recognition and specifically that key residues mutated in cancer patients impair nucleosome binding.

**Figure 4.**
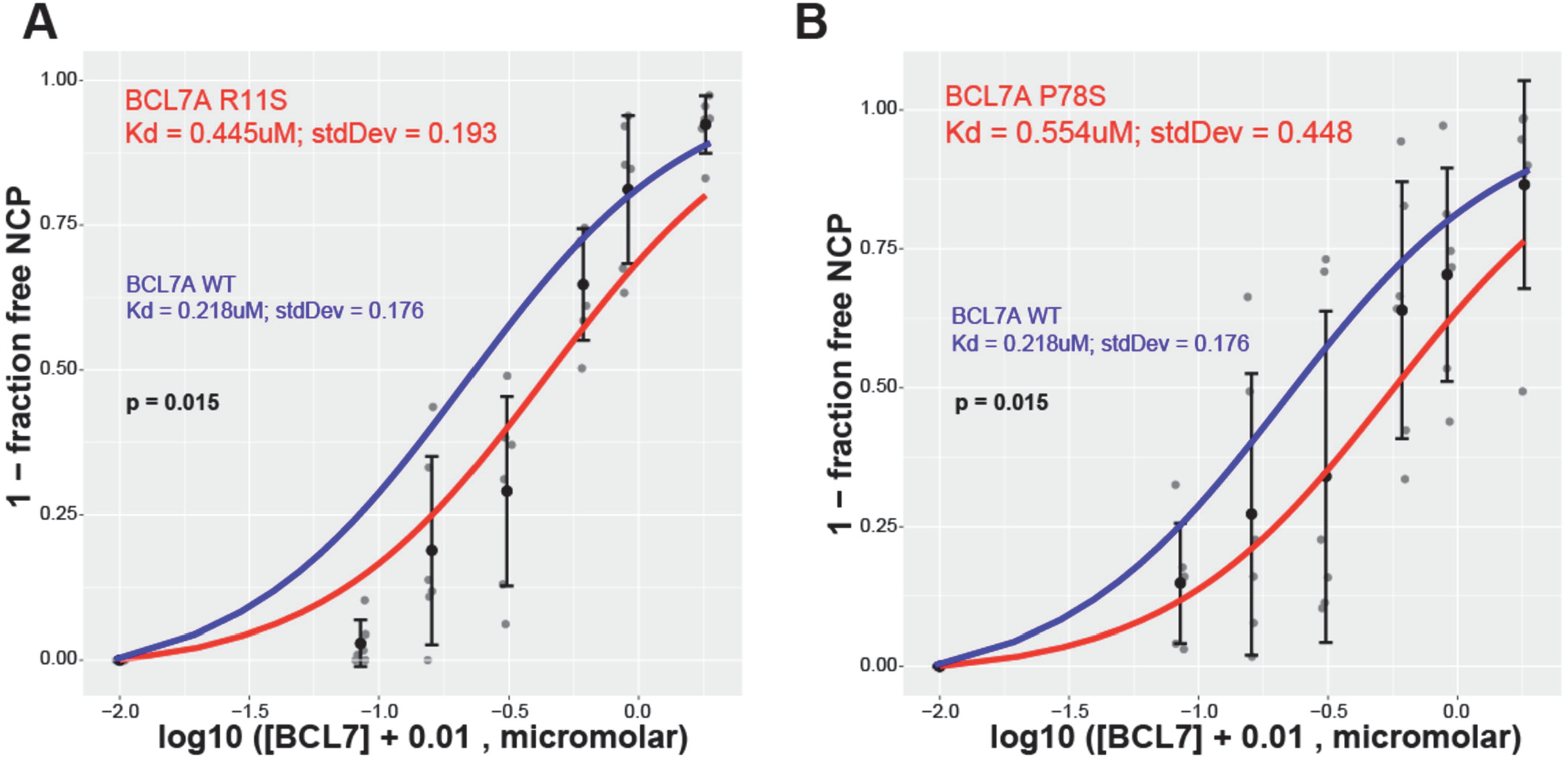
Nucleosome binding by BCL7A is impaired by cancer-derived mutations. **A)** Quantification of EMSA experiments performed with NCP probe and BCL7A WT or R11S mutant protein. The WT data (blue curve fit) is the same as shown in Fig. E2. **B)** EMSA of BCL7A P78S mutant protein binding to labelled NCP probe. The indicated p values are for one-tailed Student’s T-test, unpaired, equal variance.

### Characterization of the BCL7A-NCP complex by size-exclusion chromatography and multi-angle light scattering

The complexes formed by BCL7A and the nucleosome were analyzed by size exclusion chromatography (SEC). Nucleosomes were incubated with a 3:1 molar excess of full length BCL7A or truncated BCL7A(1-100). The samples were injected individually on a Superose-6 Increase 3.2/300 size exclusion column. Each chromatogram displays a single symmetric peak, indicating the homogeneity of the preparations. Samples containing either full-length BCL7A or BCL7A(1-100) or NCP alone were used as a control. The integrity and the composition of the complexes were assessed by running the peak fractions of the SEC purifications on native gels and SDS-PAGE gels (Fig. 5). The SEC purifications showed the formation of stable complexes for both full-length BCL7A and BCL7A(1-100) with the NCP. Interestingly, full-length BCL7A eluted in a peak corresponding to a higher molecular weight than a regular 25 kDa protein, possibly due to limited globular folding of BCL7 proteins that are characterized by a long, disordered tail.

**Figure 5.**
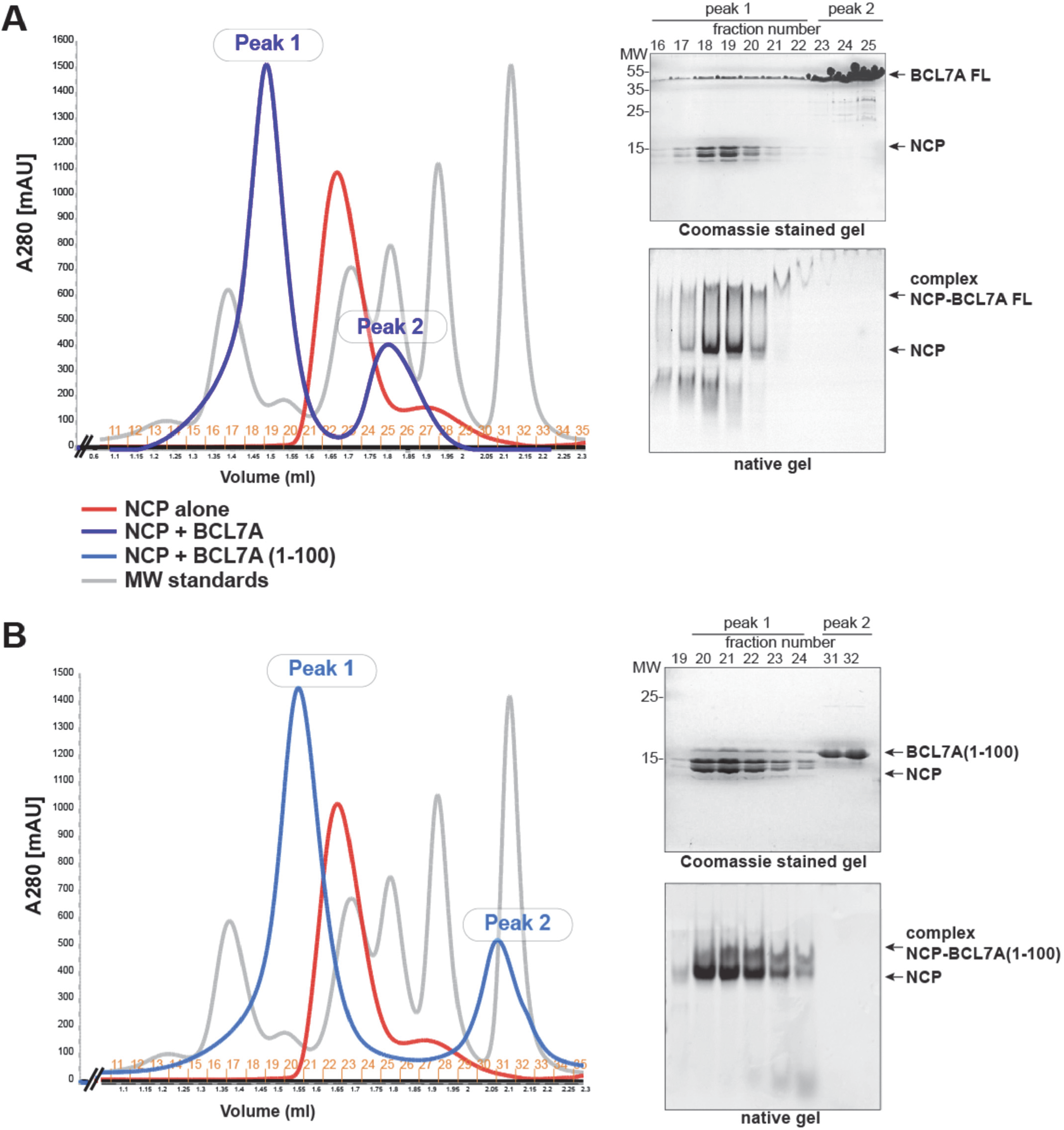
Size exclusion chromatography of the BCL7A-NCP complex. **A)** BCL7A forms a stable complex with the nucleosome core particle (NCP). Size exclusion chromatography of BCL7A bound to the NCP (3:1 molar excess of BCL7A) performed on a Superose 6 Increase column showing shorter elution time of the BCL7A-NCP complex. A_280_ traces from SEC runs are shown as colored solid lines: blue line is the BCL7A-NCP complex (peak 1) and the excess of BCL7A (peak 2), red line is the NCP alone, grey line represents standards. Top right panel shows the Coomassie stained SDS-PAGE gel of the SEC peak fractions; BCL7A and histones are indicated. Bottom right panel is an EtBr-stained native gel of the same fractions showing the integrity of the nucleosomes and a shift corresponding to complex formation is indicated. **B)** BCL7A(1-100) forms a stable complex with the nucleosome core particle (NCP). Size exclusion chromatography of BCL7A(1-100) bound to the NCP (3:1 molar excess of BCL7A(1-100)) performed on a Superose 6 Increase showing shorter elution time of the BCL7A(1-100)-NCP complex. A_280_ traces from SEC runs are shown as colored solid lines: light blue line is the BCL7A(1-100)-NCP complex (peak 1) and the excess of BCL7A(1-100) (peak 2), red line is the NCP alone, grey line represents standards. Top right panel shows the Coomassie stained SDS-PAGE gel of the SEC peak fractions; BCL7A(1-100) and histones are indicated. Bottom right panel is an EtBr-stained native gel of the same fractions showing the integrity of the nucleosomes and a shift corresponding to complex formation is indicated.

The stoichiometry of the nucleosome-BCL7A complex was investigated using SEC followed by multi-angle light scattering (MALS). The NCP alone or the complex NCP-BCL7A were injected on a Superose-6 Increase 10/300 SEC column connected to a light scattering detector. The analysis of the molar mass at the center part of the peak reveals mono-dispersity for all samples (Fig. 6). The NCP alone eluted in a single peak, with a measured molecular mass of 214.7 kDa (198.8 kDa calculated). A mixture of NCP and BCL7A, with BCL7A in large molar excess, eluted in two peaks. The first peak contained both the NCP and BCL7A, and the second peak contained BCL7A only. MALS analysis shows that the second peak corresponding to BCL7A alone has a molecular mass of 29.5 kDa (25.0 kDa calculated), indicative of a monomeric species. The first peak yields a molecular mass of 233.5 KDa (223.8 kDa calculated) which hints to the molecular mass of a 1:1 complex.

**Figure 6.**
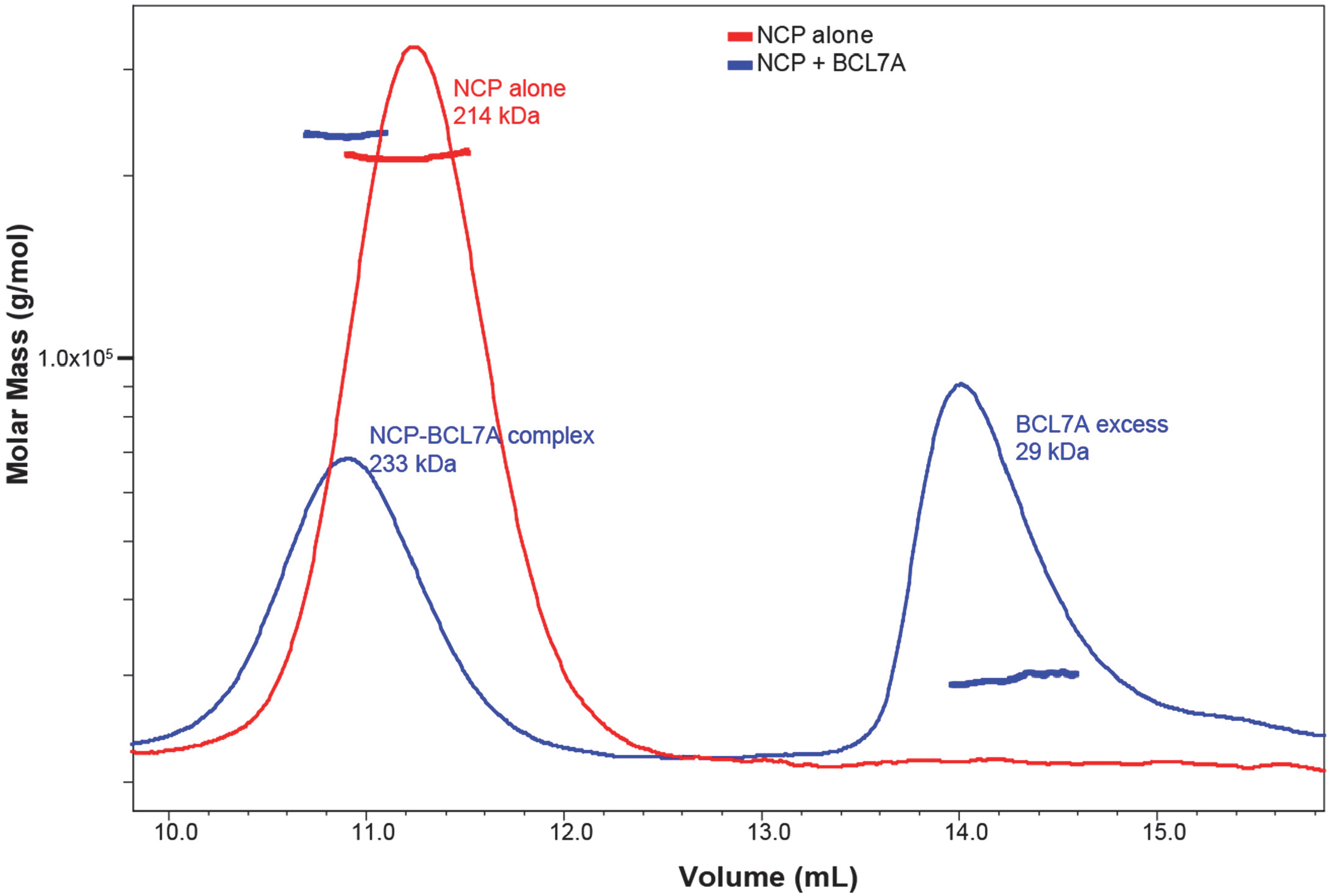
SEC-MALS analysis of nucleosomes and NCP-BCL7A complex. A_280_ traces from the SEC runs are shown as colored solid lines, plotted on the same relative scale. The MALS-derived molecular mass distributions are plotted as individual points in the colors corresponding to the A_280_ traces, with the scale shown on the left side. The samples analyzed were NCP only (red) and NCP-BCL7A complex (blue). The observed molecular mass of each sample is indicated in the respective colors. The complex between nucleosome and BCL7A occurs at a 1:1 ratio.

### Characterization of the BCL7A-NCP complex by analytical ultra-centrifugation

To better understand the NCP-BCL7A complex identified by EMSA experiments, we employed sedimentation velocity analytical ultracentrifugation (SV-AUC). Full length BCL7A is a monomer in solution with a sedimentation coefficient of 1.87 S, which corresponds to the MW of about 26 KDa, and has a high frictional coefficient due to the protein’s disordered nature (Fig. 7). The NCP alone sediments as a homogeneous 11.35 S species, consistent with earlier studies of isolated nucleosome particles^38,23^ and the complex NCP-BCL7A sediments as 10.52 S species, suggesting the formation of a 1:1 molar ratio complex. The fact that the complex NCP-BCL7A sediments more slowly than the nucleosome itself is imputable to the increased frictional coefficient of the complex compared to NCP alone due to the binding of the nucleosome to an intrinsically disordered protein as previously described^23^. Integrity of the complex NCP-BCL7A after the SV-AUC run was also confirmed by running the sample on SDS-PAGE gel (Fig S3).

**Figure 7.**
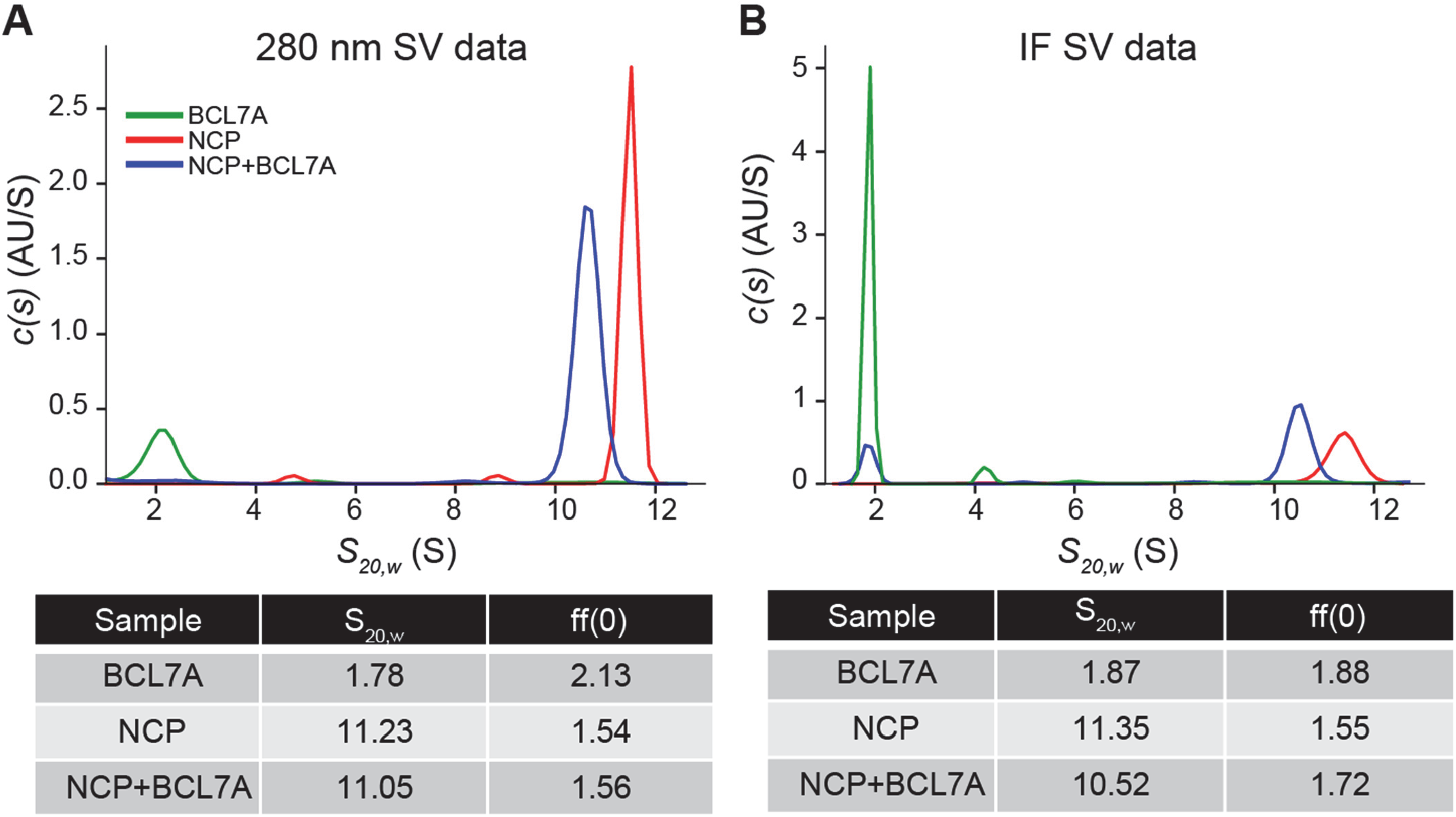
Stable complex formation between BCL7A and the nucleosome. Analytical ultracentrifugation was performed with recombinant BCL7A and NCP with a 1:5 molar excess of BCL7A. Interference sedimentation coefficient c(s) distribution profiles of NCP, NCP+BLCL7A and BCL7A samples are reported. Interference sedimentation velocity (IF SV) data were analyzed using the sedimentation coefficient distributions c(s) model and then loaded into the software GUSSI for superimposed plotting and peak integration.

## DISCUSSION

BCL7 proteins are intrinsically disordered proteins for which only limited structural and functional information exist. Here we present evidence that BCL7A binds the nucleosome with high affinity and a 1:1 stoichiometry and we identify the conserved and ordered N-terminal part of BCL7A as sufficient to nucleosome binding. We also report that BCL7 proteins bind DNA with no specificity under our experimental conditions.

The current literature contains reports of five mSWI/SNF subunits able to interact with DNA directly: the two ATPases BRG1 and BRM (SMARCA4 and SMARCA2), BAF47 (SMARCB1), BAF57 (SMARCE1), and BAF250A and BAF250B (ARID1A and ARID1B). Here we provide evidence that the BCL7 proteins belong to this list as they bind the nucleosomes and free DNA with relatively high affinity. Using EMSA, we arrived at Kd value of approximately 220 nM for the binding of BCL7A to both ligands. The results were validated using a complementary technique, MST. This binding constant is comparable to what has been reported for the bromodomain of BRG1 and BRM (around 400-600 nM, by nuclear magnetic resonance) and stronger than the bromodomain-AT-hook regions of these two proteins (16-40 µM, by fluorescence anisotropy)^39^. Another study arrived at a Kd of 2-7 µM, by isothermal titration calorimetry, for the AT-hook domain of BRG1^40^. The high-mobility-group domain of BAF57 was shown to bind specifically to cruciform (4-way junction) DNA^41^, and a Kd value of 296 nM by fluorescence quenching assay has been reported for this interaction^42^. Binding of the ARID domain of ARID1A to naked DNA occurs with a Kd of 92 nM ^43^. BAF47 binds to DNA through its N-terminal winged-helix domain with a dissociation constant estimated by NMR to be in the high micromolar range^44^. It has been proposed that several NCP- and DNA-binding activities are involved in the recruitment of the mSWI/SNF complex to chromatin^45,46^, and our work shows that BCL7 proteins are likely contributors to this process.

We found that BCL7A binds to DNA with no apparent sequence specificity, which is a characteristic shared with the other subunits mentioned above. AT-hook domains were initially reported as favoring AT-rich sequences, but later reports found no such preference for the two ATPases and the ARID proteins^47,48,49^. This apparent lack of sequence specificity is unsurprising, considering the large variety of DNA sequences the mSWI/SNF complex must be able to interact with as an important regulator of genome compaction. As consequence, we postulate that binding specificity of BCL7 proteins to the chromatin fibre is likely driven by putative contacts with the histones, with sequence-specific DNA binding proteins or through other SWI/SNF subunits, or a combination of these factors. Additionally, we cannot rule out the possibility that BCL7 proteins favor binding to some special DNA conformations that were not tested in this study (e.g. cruciform or G-quadruplexes), as has been reported for other mSWI/SNF subunits^50,51^.

Our results indicate that BCL7A binds to DNA and the NCP with comparable affinities. We could not find a report about other mSWI/SNF subunits where binding to NCP and naked DNA templates were compared using the same techniques and reagents, so direct comparisons are difficult to make. In that sense, our study appears unique in its characterization of the two binding activities of human BCL7A. However, a study of the SWARM domain of budding yeast Swi3, the homolog of human BAF155 and BAF170 (SMARCC1 and SMARCC2), reported comparable dissociation constants for naked DNA and NCPs, roughly estimated at 62 nM by EMSA^52^.

Unlike several of the other mSWI/SNF subunits mentioned above, the BCL7 proteins do not possess a discernible DNA binding domain. However, all three share a highly related N-terminal sequence of about 50 amino acids that is followed by an extended tail predicted to be disordered. A truncation experiment was performed and revealed that the N-terminal 100 amino acids of BCL7A are sufficient for NCP binding albeit with lower apparent affinity. Efforts to produce additional mutants, to better define the region of DNA binding activity, were unsuccessful in producing proteins that would behave well *in vitro*. Further experimentation will be required to determine if the C-terminal segment of the protein can also contribute to the interaction with DNA and/or the NCP, to the remodelling activity of the mSWI/SNF complex or the targeting of the complex to specific genomic sites.

Most of the biochemical work presented herein was conducted with the BCL7A protein. We also presented EMSA results showing that BCL7C also binds to the NCP with high affinity. Similar experiments were also conducted with BCL7B and indicated a similar molecular function (Fig. S4). We conclude from this that DNA and NCP binding is a property shared by all three members of the BCL7 family and we postulate that this involves the conserved N-terminus of the respective proteins. The role played by the divergent C-termini, predicted to be disordered, is less clear. It is interesting to note that although the three human proteins diverge in their C-terminal sequences, each respective protein has a C-terminus that is better conserved throughout evolution. For example, although the C-termini of human BCL7A and BCL7C have only 18 % amino acid identity past position 51, the C-termini of human and avian (*G. gallus*) BCL7A are 70 % identical. Phylogenetic conservation of these protein regions is likely due to selective pressure, implying functional constraints that may guide the elucidation of the molecular function of these protein regions.

The EMSA results show that BCL7 proteins directly bind the NCP with high affinity. The complex BCL7A-NCP was further characterized using different approaches. SEC-MALS and AUC experiments provided evidence that BCL7A forms a well-defined 1:1 complex with the nucleosome. Thus, although BCL7A exhibits features of an intrinsically disordered and extended protein it seems yet to precisely bind the nucleosome.

Keeping in consideration the SEC profiles of the BCL7A-NCP complex, the Kd calculated by the EMSA experiments, we argue that BCL7A N-terminus contributes to NCP binding. Further studies await to be conducted to investigate the role of the non-ordered region of BCL7 proteins in chromatin interactions and remodelling. The high conformational flexibility predicted for the BCL7 C-terminus probably allows molecular mechanisms that are unlikely for ordered proteins. BCL7 proteins functionality most likely requires at least some degree of conformational flexibility and structural dynamics.

The mSWI/SNF is interesting from a medical perspective because one or more of its components is disrupted in human diseases. For instance, BCL7A is the target of driver mutations in diffuse large B-cell lymphoma^17^ and is considered to function as a tumor suppressor^11^. However, the impact of these mutations on mSWI/SNF function and on the cancerous phenotype is poorly characterized. We biochemically evaluated the impact on nucleosome recognition of two missense mutations reported in patients affected by different cancers: R11S and P78S, both located in the N-terminal part of BCL7A. The protein mutants were well-behaved according to SEC-MALS analysis (data not shown), indicating that the mutations do not affect the structural integrity of BCL7 proteins. Using EMSA experiments, we calculated the binding affinity of these protein mutants and found that the mutations decrease binding affinity for the nucleosome by about 50%. The number of reported patients affected by haematological malignancy (diffuse large B cell lymphoma [DLBCL]) carrying the BCL7A R11 mutation^53,54,55^ increased significantly in the recent years and it is now considered a mutational hotspot. R11 is particularly interesting as it is located in the conserved, structured region of BCL7A, and is part of a predicted α-helix. Further, cross-linking mass spectrometry experiments^56^ have shown BS3-mediated cross-links between BCL7A or BCL7C (at K15 or K19) and several residues on αC of H2B bordering the acidic patch (between Lys109 and Lys121; coordinates based on human H2B counting the initiating Met) and with two lysines at the N-terminus of H2A.

Whether R11 is implicated in histone and/or DNA contacts remains to be established. An interesting possibility is that R11 could function as a canonical arginine (arginine anchor^57^) that inserts into the narrow cavity defined by α2 and α3 helices of H2A and the C-terminal helix of H2B, which form the nucleosome acidic patch. Structural data will be required to explore this hypothesis.

It has been reported that R11 in BCL7B is a site of the post translational modification (PTM) poly-ADP-ribosylation (PARylation)^58^. This is especially interesting because PTMs are thought to regulate interaction of chromatin proteins with the nucleosome^59^. This suggests the possibility that PARylation or other PTMs may somehow regulate association/dissociation of BCL7 and the nucleosome.

BCL7A P78 mutant was selected for its reported frequency in patients affected by lung adenocarcinoma^60^ and vicinity to the N-terminal ordered part of the molecule. Besides the acidic patch, the other nucleosomes regions that are critical for chromatin proteins binding are C-terminal helix of H2B, the histone H3 α1L1 elbow region and the histone H2B α1L1 elbow^61^. If the N-terminal part of BCL7 proteins targets the acidic patch of the nucleosome, there is the possibility that BCL7A flexible region encompassing P78 could arrange to become in contact with the other regions of the nucleosome or could make contact with nucleosomal DNA.

Our biochemical studies provide a molecular basis for understanding the deleterious effects of some missense mutations in BCL7 proteins on binding to chromatin. Structural information on BCL7 proteins would allow mapping of these mutations on the structure and provide further mechanistic insight on the role of the mutations on BCL7 function.

## Acknowledgements

This research was funded in part by the Fondation pour la Recherche Médicale (grant number AJE20171039021), an ATIP-Avenir award (contract # 188435), an award from La Ligue Contre le Cancer (JMG/SP - n° 01B.2021), startup funds from the IGBMC, and salary support by the Centre National de la Recherche Scientifique to E.B. D.D. was partially supported by a 4^th^ year fellowship from the Fondation pour la Recherche Médicale. J.L. is supported by a scholarship from the École Doctorale de l’Université de Strasbourg. A.B. is supported by an operating grant from the Canadian Institutes of Health Research (PJT-183839). The authors wish to thank Irwin Davidson for support and helpful discussions, the staff at the Centre de Biologie Intégrative of the IGBMC structural biology platform for technical assistance, and Denis Fumagalli from the Mediaprep platform.

## FIGURES

**Figure S1.**
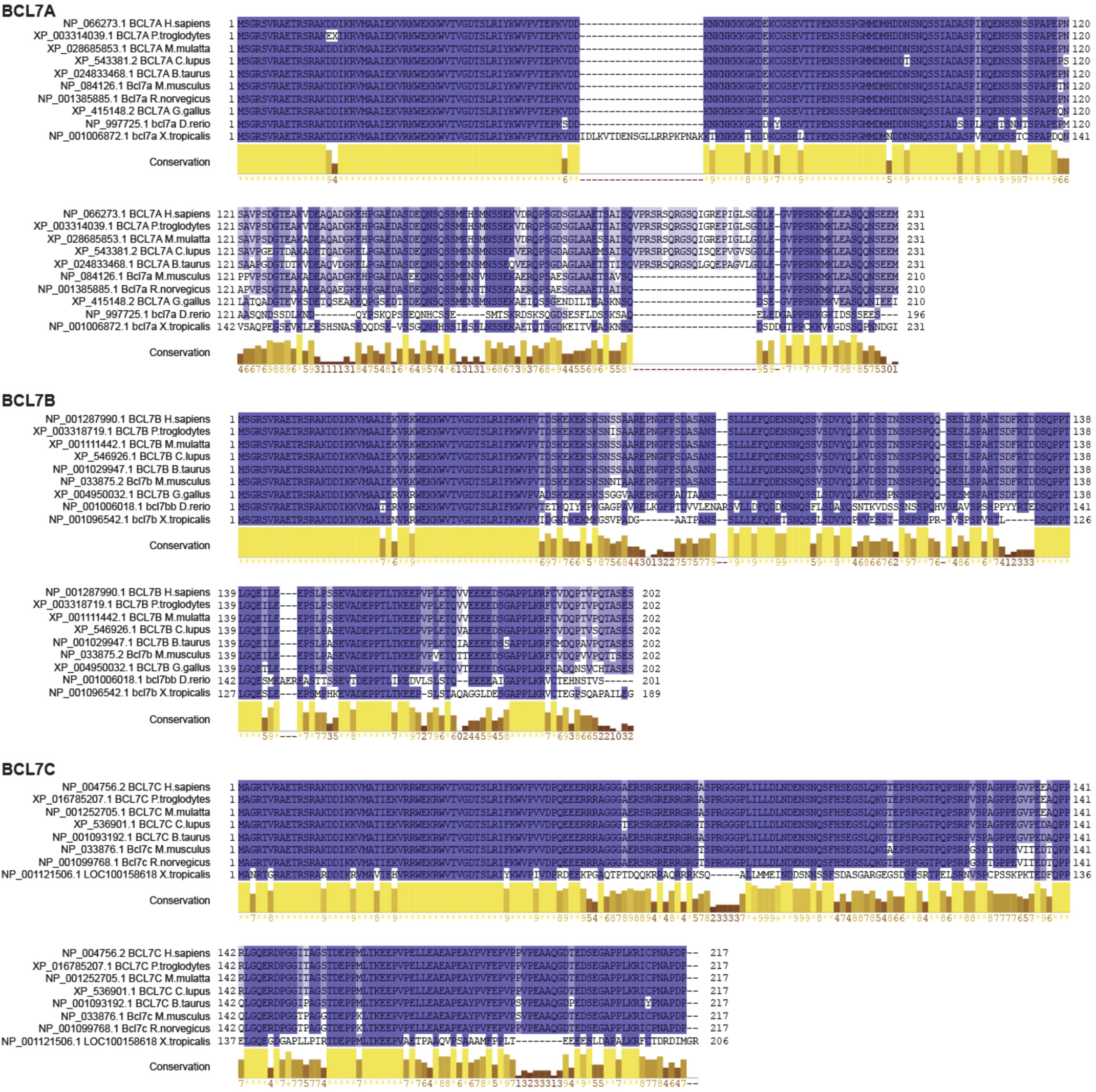
Phylogenetic conservation of the three BCL7 proteins. Multiple sequence alignment using JalView and the Muscle algorithm for the BCL7 protein sequences from diverse species. The species shown are those reported by the NCBI Homologene website as the most closely related. Coloring is by percentage identity of sequence.

**Figure S2.**
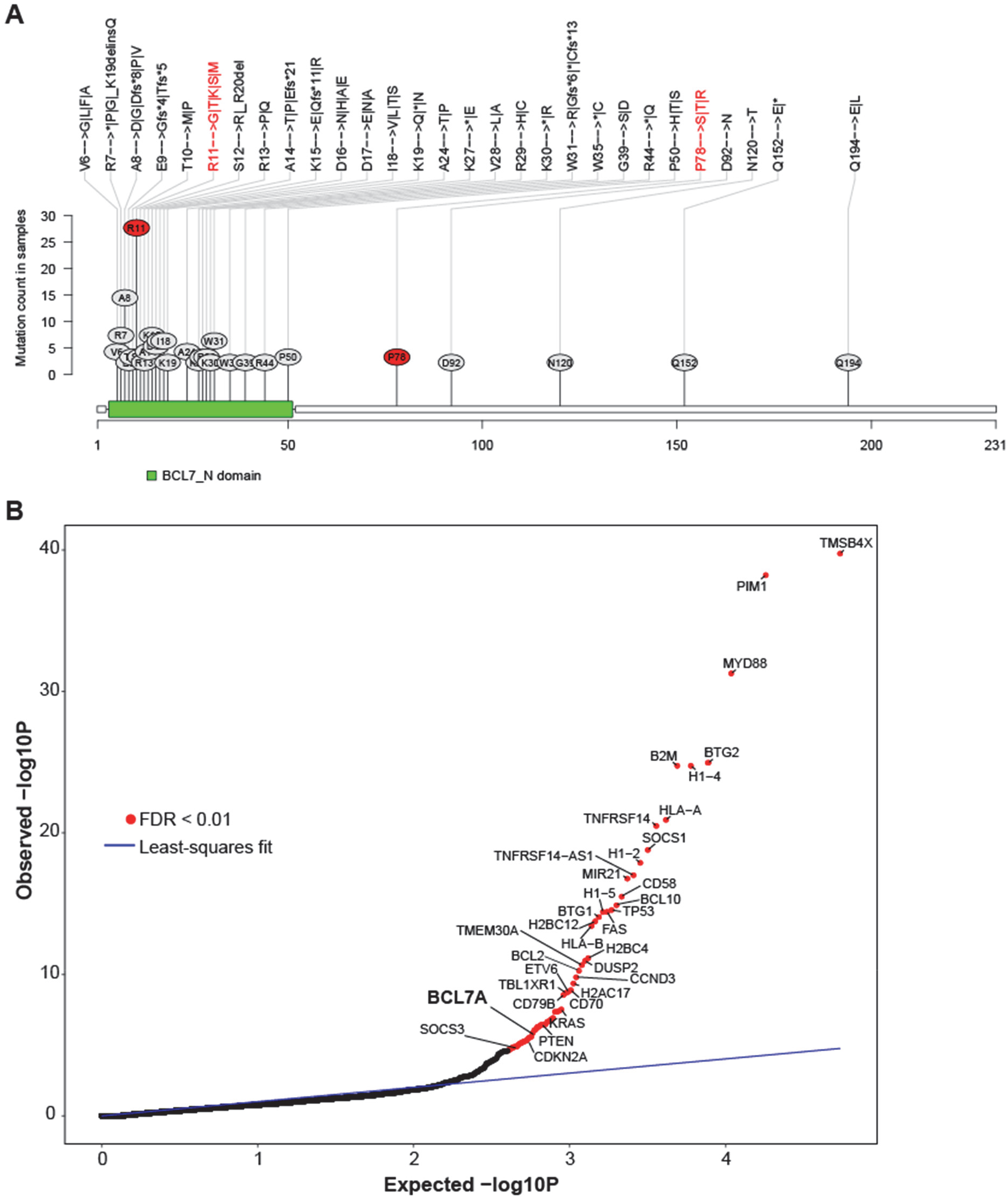
Cancer-associated mutations of the BCL7A gene. **A)** Mutational data regarding the BCL7A gene from the COSMIC database. For clarity, only amino acid residues affected in two or more samples (database entries) are represented. Labels at the top indicate mutation results at the protein level; changes are listed in decreasing order of frequency (e.g. the R11G mutation is more frequent than R11M). Residues in red were tested in EMSA experiments. **B)** Quantile-quantile plot of observed *versus* expected p-values of hotspot mutations in DLBCL. Genes with an FDR value less than 0.01 (Benjamini-Hochberg algorithm on the observed p-values) are considered significantly affected by driver mutations. The blue line represents the least-square fit on the quantile-quantile data, forcing the intercept at zero. The BCL7A gene (FDR = 8.10E-04) ranks with other well-known tumor suppressors and proto-oncogenes.

**Figure S3.**
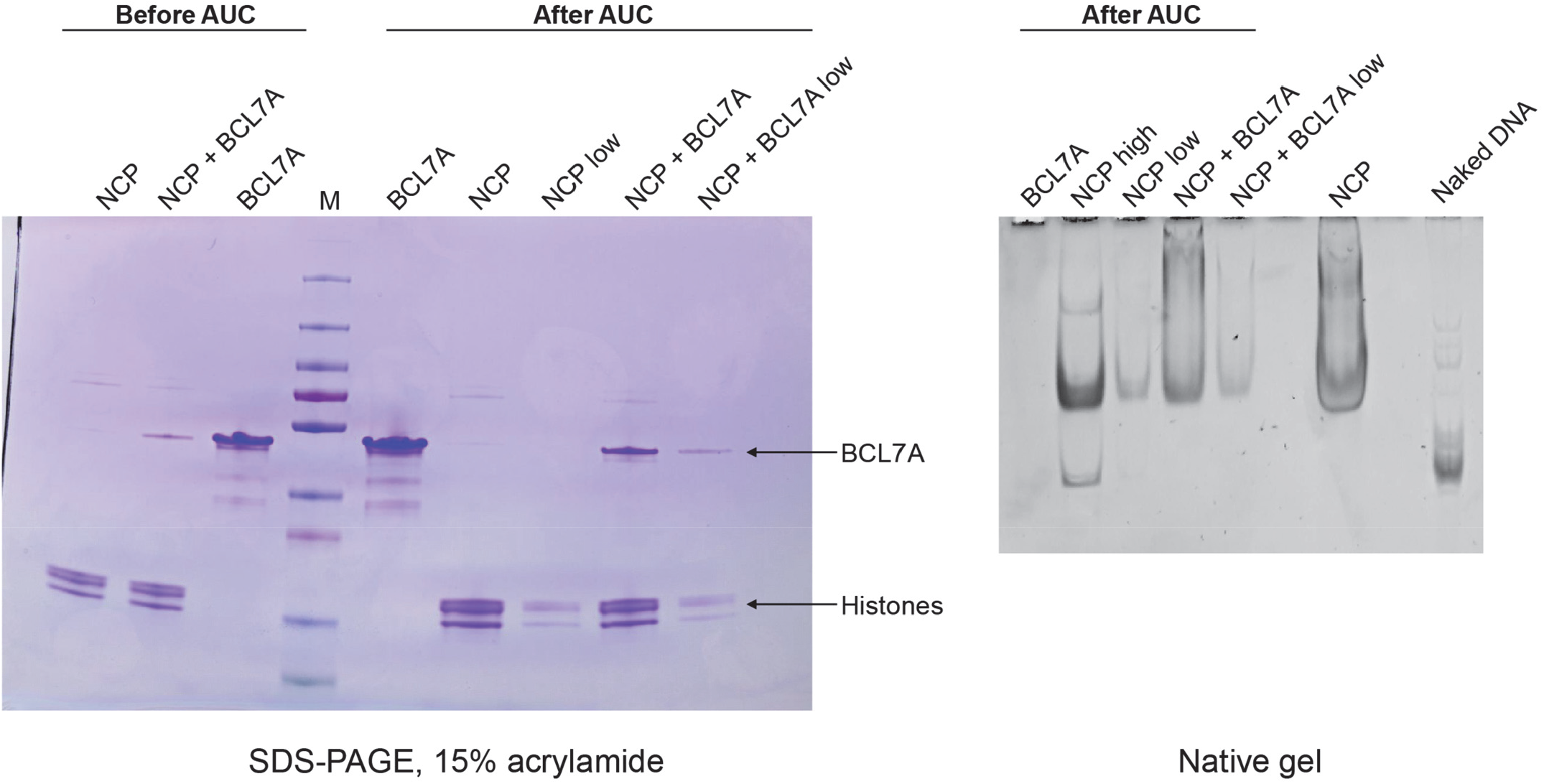
Analysis of samples before and after AUC run. Protein samples were analyzed by SDS-PAGE followed by Coomassie blue staining (left) or by native gel EMSA followed by ethidium bromide staining (right). Samples before or after the centrifugation are shown. “Low” indicates that the AUC run was performed using sample components at a lower concentration.

**Figure S4.**
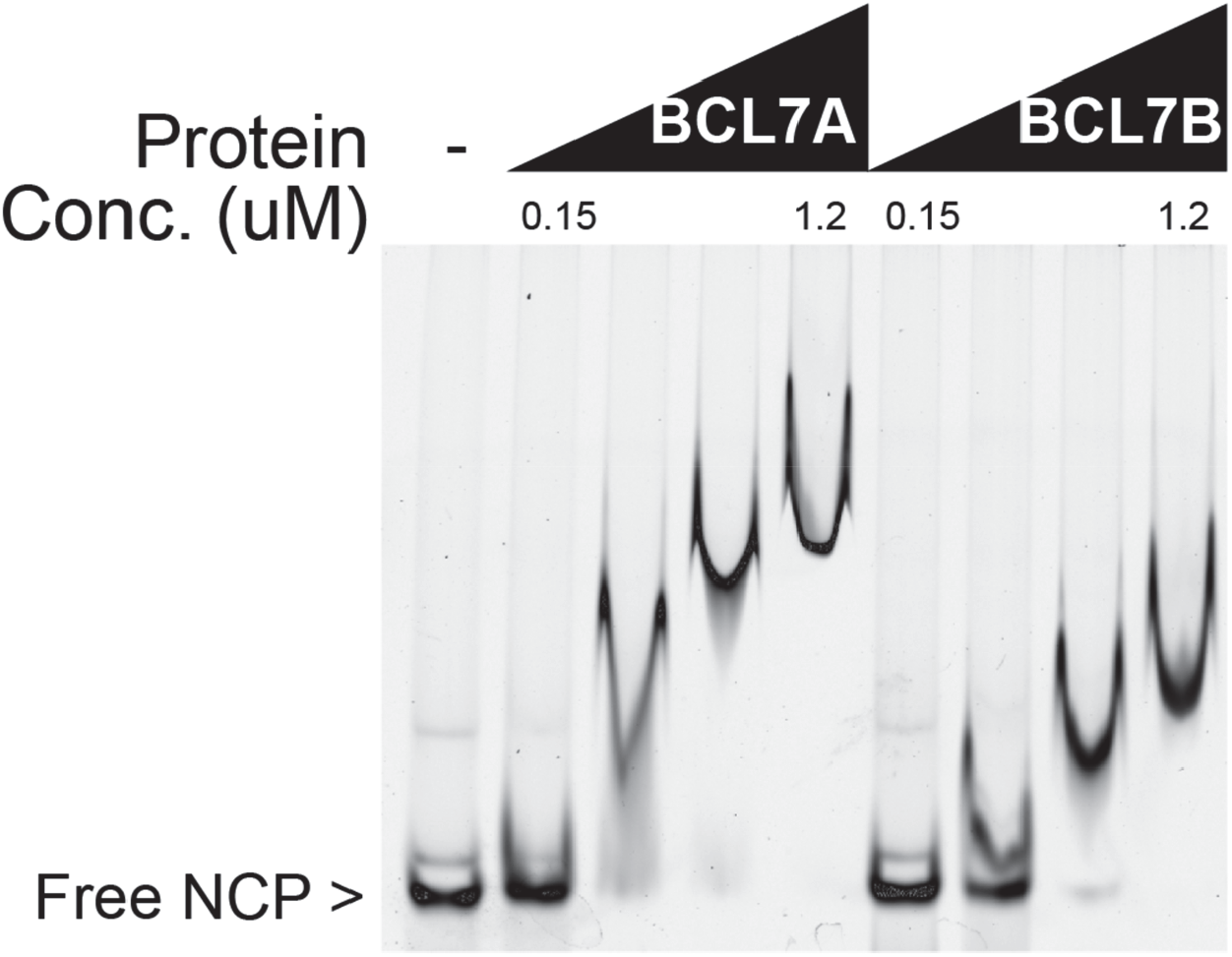
NCP binding by BCL7B. EMSA performed on Cy5-labelled NCP using increasing amounts of purified BCL7A or BCL7B proteins.

---

**Table S1.**
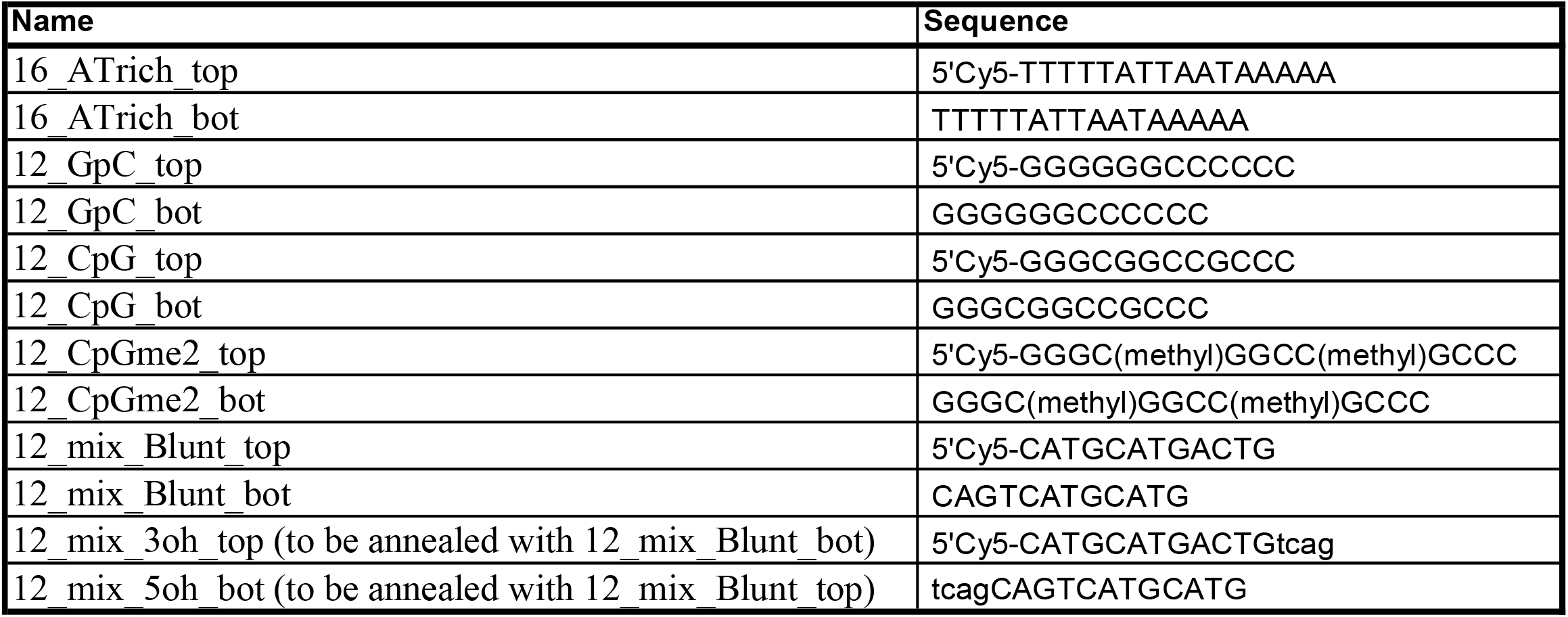
List of oligonucleotides used in EMSA experiments

## REFERENCES

1. Clapier, C. R. & Cairns, B. R. The biology of chromatin remodeling complexes. Annu. Rev. Biochem. 78, 273–304 (2009).

2. Narlikar, G. J., Sundaramoorthy, R. & Owen-Hughes, T. Mechanisms and functions of ATP-dependent chromatin-remodeling enzymes. Cell 154, 490–503 (2013).

3. Middeljans, E. et al. SS18 Together with Animal-Specific Factors Defines Human BAF-Type SWI/SNF Complexes. PLoS ONE 7, e33834 (2012).

4. Kadoch, C. et al. Proteomic and bioinformatic analysis of mammalian SWI/SNF complexes identifies extensive roles in human malignancy. Nat. Genet. 45, 592–601 (2013).

5. Marcon, E. et al. Human-Chromatin-Related Protein Interactions Identify a Demethylase Complex Required for Chromosome Segregation. Cell Rep. 8, 297–310 (2014).

6. Huttlin, E. L. et al. The BioPlex Network: A Systematic Exploration of the Human Interactome. Cell 162, 425–440 (2015).

7. Wan, C. et al. Panorama of ancient metazoan macromolecular complexes. Nature 525, 339–344 (2015).

8. Siguero-Álvarez, M. et al. A Human Hereditary Cardiomyopathy Shares a Genetic Substrate With Bicuspid Aortic Valve. Circulation 147, 47–65 (2023).

9. Wischhof, L. et al. BCL7A -containing SWI/SNF/BAF complexes modulate mitochondrial bioenergetics during neural progenitor differentiation. EMBO J. 41, p(2022).

10. Huang, C., Hao, Q., Shi, G., Zhou, X. & Zhang, Y. BCL7C suppresses ovarian cancer growth by inactivating mutant p53. J. Mol. Cell Biol. 13, 141–150 (2021).

11. Baliñas-Gavira, C. et al. Frequent mutations in the amino-terminal domain of BCL7A impair its tumor suppressor role in DLBCL. Leukemia 34, 2722–2735 (2020).

12. Wischhof, L. et al. The SWI/SNF subunit Bcl7a contributes to motor coordination and Purkinje cell function. Sci. Rep. 7, 17055 (2017).

13. Uehara, T., Kage-Nakadai, E., Yoshina, S., Imae, R. & Mitani, S. The Tumor Suppressor BCL7B Functions in the Wnt Signaling Pathway. PLoS Genet. 11, e1004921 (2015).

14. Kadoch, C. et al. Proteomic and bioinformatic analysis of mammalian SWI/SNF complexes identifies extensive roles in human malignancy. Nat. Genet. 45, 592–601 (2013).

15. Izhar, L. et al. A Systematic Analysis of Factors Localized to Damaged Chromatin Reveals PARP-Dependent Recruitment of Transcription Factors. Cell Rep. 11, 1486–1500 (2015).

16. Forbes, S. A. et al. COSMIC: exploring the world’s knowledge of somatic mutations in human cancer. Nucleic Acids Res. 43, D805–D811 (2015).

17. Reddy, A. et al. Genetic and Functional Drivers of Diffuse Large B Cell Lymphoma. Cell 171, 481–494.e15 (2017).

18. Mashtalir, N. et al. Modular Organization and Assembly of SWI/SNF Family Chromatin Remodeling Complexes. Cell 175, 1272–1288.e20 (2018).

19. Wang, L. et al. Structure of nucleosome-bound human PBAF complex. Nat. Commun. 13, 7644 (2022).

20. He, S. et al. Structure of nucleosome-bound human BAF complex. Science 367, 875–881 (2020).

21. Wagner, F. R. et al. Structure of SWI/SNF chromatin remodeller RSC bound to a nucleosome. Nature 579, 448–451 (2020).

22. Han, Y., Reyes, A. A., Malik, S. & He, Y. Cryo-EM structure of SWI/SNF complex bound to a nucleosome. Nature 579, 452–455 (2020).

23. Yang, C., van der Woerd, M. J., Muthurajan, U. M., Hansen, J. C. & Luger, K. Biophysical analysis and small-angle X-ray scattering-derived structures of MeCP2-nucleosome complexes. Nucleic Acids Res. 39, 4122–4135 (2011).

24. Mashtalir, N. et al. A Structural Model of the Endogenous Human BAF Complex Informs Disease Mechanisms. Cell 183, 802–817.e24 (2020).

25. Liu, X., Li, M., Xia, X., Li, X. & Chen, Z. Mechanism of chromatin remodelling revealed by the Snf2-nucleosome structure. Nature 544, 440–445 (2017).

26. Valencia, A. M. et al. Recurrent SMARCB1 Mutations Reveal a Nucleosome Acidic Patch Interaction Site That Potentiates mSWI/SNF Complex Chromatin Remodeling. Cell 179, 1342–1356.e23 (2019).

27. Phelan, M. L., Sif, S., Narlikar, G. J. & Kingston, R. E. Reconstitution of a Core Chromatin Remodeling Complex from SWI/SNF Subunits. Mol. Cell 3, 247–253 (1999).

28. Lowary, P. T. & Widom, J. New DNA sequence rules for high affinity binding to histone octamer and sequence-directed nucleosome positioning. J. Mol. Biol. 276, 19–42 (1998).

29. Dyer, P. N. et al. Reconstitution of Nucleosome Core Particles from Recombinant Histones and DNA. in Methods in Enzymology vol. 375 23–44 (Elsevier, 2003).

30. Shim, Y., Duan, M.-R., Chen, X., Smerdon, M. J. & Min, J.-H. Polycistronic coexpression and nondenaturing purification of histone octamers. Anal. Biochem. 427, 190–192 (2012).

31. Jarmoskaite, I., AlSadhan, I., Vaidyanathan, P. P. & Herschlag, D. How to measure and evaluate binding affinities. eLife 9, e57264 (2020).

32. Schuck, P. Size-Distribution Analysis of Macromolecules by Sedimentation Velocity Ultracentrifugation and Lamm Equation Modeling. Biophys. J. 78, 1606–1619 (2000).

33. Brautigam, C. A. Calculations and Publication-Quality Illustrations for Analytical Ultracentrifugation Data. Methods Enzym. 562, 109–133 (2015).

34. Mlynarczyk, C. et al. BTG1 mutation yields supercompetitive B cells primed for malignant transformation. Science 379, eabj7412 (2023).

35. ICGC/TCGA Pan-Cancer Analysis of Whole Genomes Consortium. Pan-cancer analysis of whole genomes. Nature 578, 82–93 (2020).

36. Mészáros, B., Erdos, G. & Dosztányi, Z. IUPred2A: context-dependent prediction of protein disorder as a function of redox state and protein binding. Nucleic Acids Res. 46, W329–W337 (2018).

37. Varadi, M. et al. AlphaFold Protein Structure Database: massively expanding the structural coverage of protein-sequence space with high-accuracy models. Nucleic Acids Res. 50, D439–D444 (2022).

38. Bradbury, E. M. K. E. Van Holde. Chromatin. Series in molecular biology. Springer-Verlag, New York. 1988. 530 pp. $ 98.00. J. Mol. Recognit. 2, – (1989).

39. Morrison, E. A. et al. DNA binding drives the association of BRG1/hBRM bromodomains with nucleosomes. Nat. Commun. 8, 16080 (2017).

40. Singh, M., D’Silva, L. & Holak, T. A. DNA-binding properties of the recombinant high-mobility-group-like AT-hook-containing region from human BRG1 protein. Biol. Chem. 387, 1469–1478 (2006).

41. Wang, W. et al. Architectural DNA binding by a high-mobility-group/kinesin-like subunit in mammalian SWI/SNF-related complexes. Proc. Natl. Acad. Sci. U. S. A. 95, 492–498 (1998).

42. Heo, Y. et al. Crystal structure of the HMG domain of human BAF57 and its interaction with fourway junction DNA. Biochem. Biophys. Res. Commun. 533, 919–924 (2020).

43. Chandler, R. L. et al. ARID1a-DNA interactions are required for promoter occupancy by SWI/SNF. Mol. Cell. Biol. 33, 265–280 (2013).

44. Allen, M. D., Freund, S. M. V., Zinzalla, G. & Bycroft, M. The SWI/SNF Subunit INI1 Contains an NTerminal Winged Helix DNA Binding Domain that Is a Target for Mutations in Schwannomatosis. Struct. Lond. Engl. 1993 23, 1344–1349 (2015).

45. Wu, J. I., Lessard, J. & Crabtree, G. R. Understanding the words of chromatin regulation. Cell 136, 200–206 (2009).

46. Ho, P. J., Lloyd, S. M. & Bao, X. Unwinding chromatin at the right places: how BAF is targeted to specific genomic locations during development. Dev. Camb. Engl. 146, dev178780 (2019).

47. Dallas, P. B. et al. The human SWI-SNF complex protein p270 is an ARID family member with non-sequence-specific DNA binding activity. Mol. Cell. Biol. 20, 3137–3146 (2000).

48. Nie, Z. et al. A specificity and targeting subunit of a human SWI/SNF family-related chromatin-remodeling complex. Mol. Cell. Biol. 20, 8879–8888 (2000).

49. Wilsker, D. et al. The DNA-binding properties of the ARID-containing subunits of yeast and mammalian SWI/SNF complexes. Nucleic Acids Res. 32, 1345–1353 (2004).

50. Makowski, M. M. et al. Global profiling of protein-DNA and protein-nucleosome binding affinities using quantitative mass spectrometry. Nat. Commun. 9, 1653 (2018).

51. Zhang, X., Spiegel, J., Martínez Cuesta, S., Adhikari, S. & Balasubramanian, S. Chemical profiling of DNA G-quadruplex-interacting proteins in live cells. Nat. Chem. 13, 626–633 (2021).

52. Da, G. et al. Structure and function of the SWIRM domain, a conserved protein module found in chromatin regulatory complexes. Proc. Natl. Acad. Sci. U. S. A. 103, 2057–2062 (2006).

53. Lacy, S. E. et al. Targeted sequencing in DLBCL, molecular subtypes, and outcomes: a Haematological Malignancy Research Network report. Blood 135, 1759–1771 (2020).

54. Gunawardana, J. et al. Recurrent somatic mutations of PTPN1 in primary mediastinal B cell lymphoma and Hodgkin lymphoma. Nat. Genet. 46, 329–335 (2014).

55. Pillonel, V. et al. High-throughput sequencing of nodal marginal zone lymphomas identifies recurrent BRAF mutations. Leukemia 32, 2412–2426 (2018).

56. Mashtalir, N. et al. A Structural Model of the Endogenous Human BAF Complex Informs Disease Mechanisms. Cell 183, 802–817.e24 (2020).

57. McGinty, R. K., Henrici, R. C. & Tan, S. Crystal structure of the PRC1 ubiquitylation module bound to the nucleosome. Nature 514, 591–596 (2014).

58. Hendriks, I. A., Larsen, S. C. & Nielsen, M. L. An Advanced Strategy for Comprehensive Profiling of ADP-ribosylation Sites Using Mass Spectrometry-based Proteomics. Mol. Cell. Proteomics MCP 18, 1010–1026 (2019).

59. McGinty, R. K. & Tan, S. Principles of nucleosome recognition by chromatin factors and enzymes. Curr. Opin. Struct. Biol. 71, 16–26 (2021).

60. Shi, J. et al. Somatic Genomics and Clinical Features of Lung Adenocarcinoma: A Retrospective Study. PLoS Med. 13, e1002162 (2016).

61. McGinty, R. K. & Tan, S. Principles of nucleosome recognition by chromatin factors and enzymes. Curr. Opin. Struct. Biol. 71, 16–26 (2021).

